# A DNA-based optical force sensor for live-cell applications

**DOI:** 10.1101/2021.12.21.473677

**Authors:** Christina Jayachandran, Arindam Ghosh, Meenakshi Prabhune, Jonathan Bath, Andrew J. Turberfield, Lara Hauke, Jörg Enderlein, Florian Rehfeldt, Christoph F. Schmidt

**Affiliations:** Third Institute of Physics—Biophysics, Georg August University, Friedrich-Hund-Platz 1, 37077 Göttingen, Germany; Department of Pharmacy - Center for Drug Research, Pharmaceutical Biology, Ludwig-Maximilians University, Butenandtstr. 5-13, 81377 Munich, Germany; Department of Biotechnology and Biophysics, Biocenter, University of Würzburg, Am Hubland, 97074 Würzburg, Germany; Synthego Corporation, 3565 Haven Ave, Menlo Park, CA 94025, USA; Department of Physics, Clarendon Laboratory, University of Oxford, Oxford, UK, OX1 3PU; The Kavli Institute for Nanoscience Discovery, University of Oxford, Oxford, UK, OX1 3QU; Institute of Pharmacology and Toxicology, University Medical Center Göttingen, Germany; Cluster of Excellence “Multiscale Bioimaging: from Molecular Machines to Networks of Excitable Cells” (MBExC), Georg August University, 37077 Göttingen, Germany; Experimental Physics I, University of Bayreuth, Universitätsstr. 30, 95440 Bayreuth, Germany; Department of Physics and Soft Matter Center, Duke University, Durham, NC 27708, USA

## Abstract

Mechanical forces are relevant for many biological processes, from wound healing or tumour formation to cell migration and differentiation. Cytoskeletal actin is largely responsible for responding to forces and transmitting them in cells, while also maintaining cell shape and integrity. Here, we describe a novel approach to employ a FRET-based DNA force sensor *in vitro* and *in cellulo* for non-invasive optical monitoring of intracellular mechanical forces. We use fluorescence lifetime imaging to determine the FRET efficiency of the sensor, which makes the measurement robust against intensity variations. We demonstrate the applicability of the sensor by monitoring cross-linking activity in *in vitro* actin networks by bulk rheology and confocal microscopy. We further demonstrate that the sensor readily attaches to stress fibers in living cells which opens up the possibility of live-cell force measurements.

---

The actin cytoskeleton ^1^ is a main component of the dominant force-generating machinery in most cells. For example, stress fibers ^2^ produce contractile forces ^3, 4^ which help in cell locomotion, division ^5^ and differentiation ^6^. Propulsive forces of ca. 20 pN are generated by actin polymerization ^7^ driving cell migration ^8–10^. In conjunction with myosin motors, actin also serves as a key element in mechanosensation ^11, 12^. Force-transmitting and sensing structures and their movements can readily be imaged in fluorescence microscopy. For example, on stiff elastic substrates, cells form prominent actin stress fibres ^13–15^. Forces and stresses, however, are not directly visible, and it remains challenging to quantitate forces on actin structure in cells due to the lack of appropriate force sensors.

Several methods have been applied to measure cellular forces transmitted to their surroundings, including traction force microscopy ^16^, atomic force microscopy ^17, 18^, and optical ^19, 20^ and magnetic tweezers ^21^. These methods are insensitive to internally balanced forces in the cells and thus cannot fully characterize cellular stresses. Recent additions to this set of methods are genetically expressed molecular force sensors (MFS) that rely on Förster resonance energy transfer (FRET). FRET-based MFS allow one to sample stresses by measuring the energy transfer efficiency between a donor and an acceptor fluorophore (FRET pair) that are coupled via a molecular spring ^22^. These MFS offer pico-Newton (pN) sensitivity and high spatial (∼20 nm) and temporal resolution (∼ ms) while minimally perturbing the cells ^23^. However, genetically expressed sensors typically use fluorescent proteins ^24, 25^ that can unfold upon force application and that possess photophysical properties inferior to those of organic dyes. When using polymer chains as molecular springs between donor and acceptor ^26, 27^, it is difficult to precisely adjust the spring stiffness. Both these limitations can be overcome by DNA-based MFS, which can be adjusted to a broad range of physiologically relevant forces ^28^ due to the easy designability of DNA structures^29–31^ as referenced in Prabhune *et al*. ^32^. DNA-based MFS grafted on surfaces have been used to investigate interfacial forces between cells and ligands.^30, 31^

Here, we present a novel DNA-based MFS for *in cellulo* and *in vitro* applications. The structure of this sensor is a DNA hairpin that can switch between two conformational states ^33, 34^. A FRET pair consisting of an organic dye (donor) and a quencher (acceptor) provides fluorescence read-out through a change in FRET efficiency when an external force pulls the hairpin apart. The hairpin switches reversibly between two states: it opens at a specific threshold force and folds back when the force is lifted. The threshold opening force of our sensors is estimated to be ≈ 10 pN ^35, 36^. In the following sections, we present the design and fabrication of this sensor, thoroughly characterize it by fluorescence lifetime spectroscopy, explore novel attachment strategies, check its performance in reconstituted actin networks, and show that the sensors can be inserted into and targeted to the actin cytoskeleton in living cells.

## Results and Discussion

### DNA sensor design

Our DNA force sensor consists of a hairpin (Fig. 1A), with a stem (8 base pairs (bp)), a loop (16 bp) and two arms (each 20 nt). Two other strands, namely the F and Q strands, hybridize to the hairpin arms. The FRET pair is formed by the fluorophore Alexa488 attached to the F strand and a quencher molecule (Iowa black FQ) attached to the Q strand. Upon hybridization with the hairpin arms, the dye and the quencher come into FRET range. The sensor is attached to actin through LifeAct ^38^, a transient actin-binding protein (ABP) with a 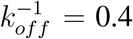 in the following way. The 5’ and 3’ termini of the hairpin sensor are modified to incorporate a HaloTag® ligand which binds covalently to the HaloTag® protein ^37^ which is expressed as a genetic fusion with red fluorescent protein (RFP) and LifeAct (Fig. 1B) (see ‘Methods’ for details). The LifeAct construct can be used for *in cellulo* measurements via transfection with the three fused genes, and it can be recombinantly expressed and purified for *in vitro* measurements. When an external force acts on this sensor and exceeds a certain threshold, the sensor hairpin undergoes a conformational switch by unfolding from a closed (high FRET) to an open (low FRET) state (Fig. 1A).

**Figure 1:**
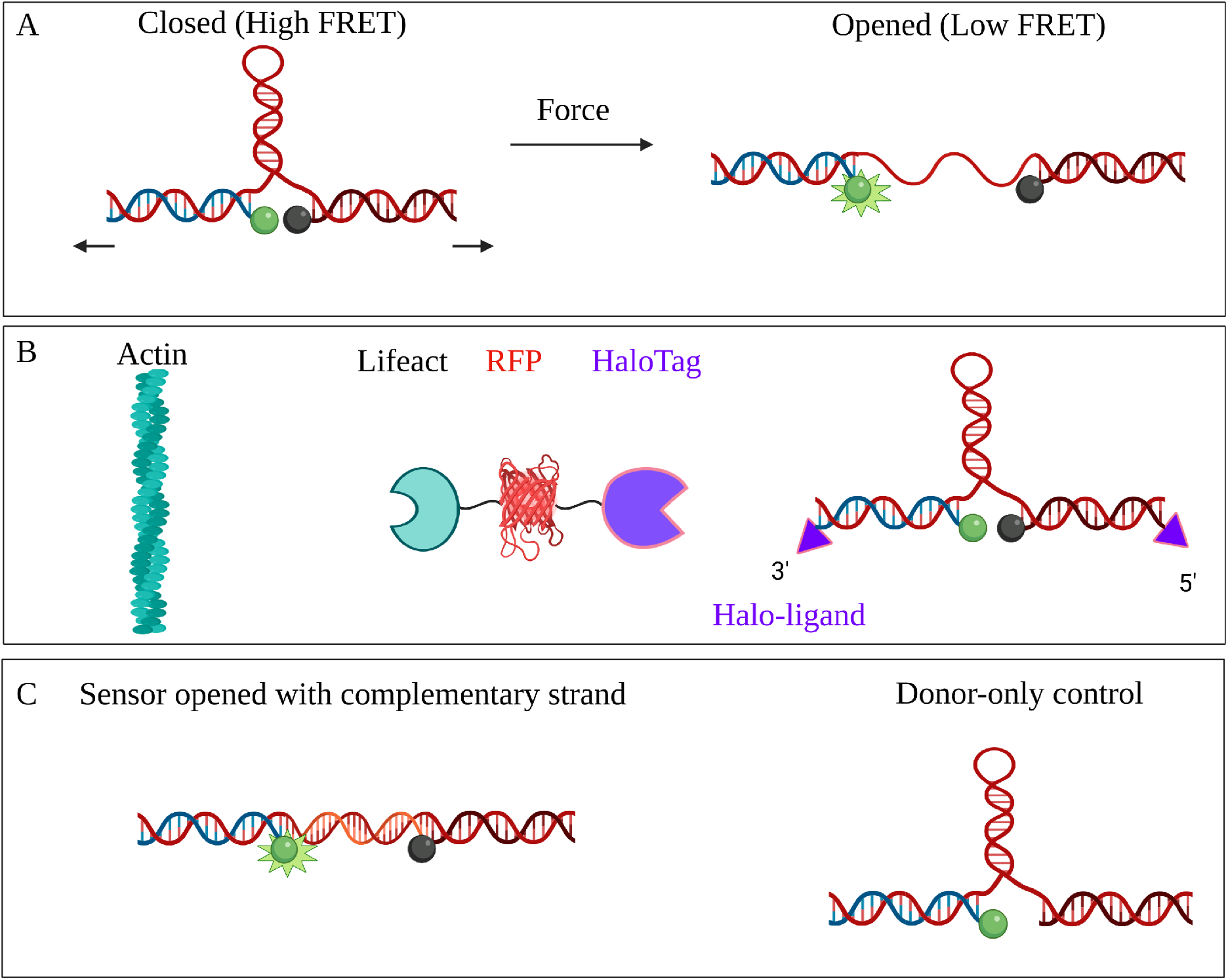
Design and attachment of the DNA force sensor. **A:** The DNA force sensor consists of a hairpin (red) with stem (8 bp), loop (16 bp) and two arms (each 20 nt). The arms hybridize to two strands, each 20 nt long, bearing the fluorophore Alexa 488 (F strand, blue) and quencher Iowa black FQ (Q strand, black) which form the FRET pair. A threshold force applied to the ends of the sensor opens it, switching it from its quenched state (high FRET) to its fluorescent state (low FRET). **B:** The outer ends of the F and Q strands are modified with a HaloTag® ligand. The HaloTag® ligands bind covalently to HaloTag®s via HaloTag® fusion ^37^. HaloTag®s were genetically expressed in the cells, as a fusion with RFP and LifeAct, an actin-binding peptide ^38^. **C:** For characterization purposes, sensor hairpins were opened using a complementary strand (orange). A control probe was designed lacking the quencher.

### Fluorescence intensity-based validation

We first characterized the sensor by bulk fluorescence intensity measurements (for experimental details see ‘Methods’). Fluorescence intensity of sensors was monitored using a standard fluorescence spectrometer (AMINCO-Bowman Series 2 Luminescence Spectrometer). A donor-only control (Fig. 1C) used as a reference to assess the FRET efficiency of the full sensor and a construct for which the hairpin loop could be opened by hybridization with a complementary DNA strand (C strand) (Fig. 1C), was used as a mimic of the sensor’s open state. In the remainder of this manuscript, we will use the following nomenclature for the various constructs: ‘closed sensors’ - sensors in their assembled geometry having both dye and quencher molecules, ‘donor-only control’ - assembled sensors that do not contain quenchers, and ‘opened sensors’ - sensors containing both fluorophore and quencher that are opened with the C strand. Closed sensors exhibit a fluorescence intensity which is ca. 15 times weaker than that of opened sensors (see Fig. 2A), confirming strong FRET-based quenching. We checked the sensor fluorescence of closed and opened sensors when cross-linked to an *in vitro* reconstituted actin network (see Fig. 2B). In this case, we observed reduced quenching for closed sensors compared to unattached sensors in solution (Fig. 2A): the fluorescence intensity of closed sensors was 50% of that of opened sensors when cross-linked to actin (Fig. 2B).

**Figure 2:**
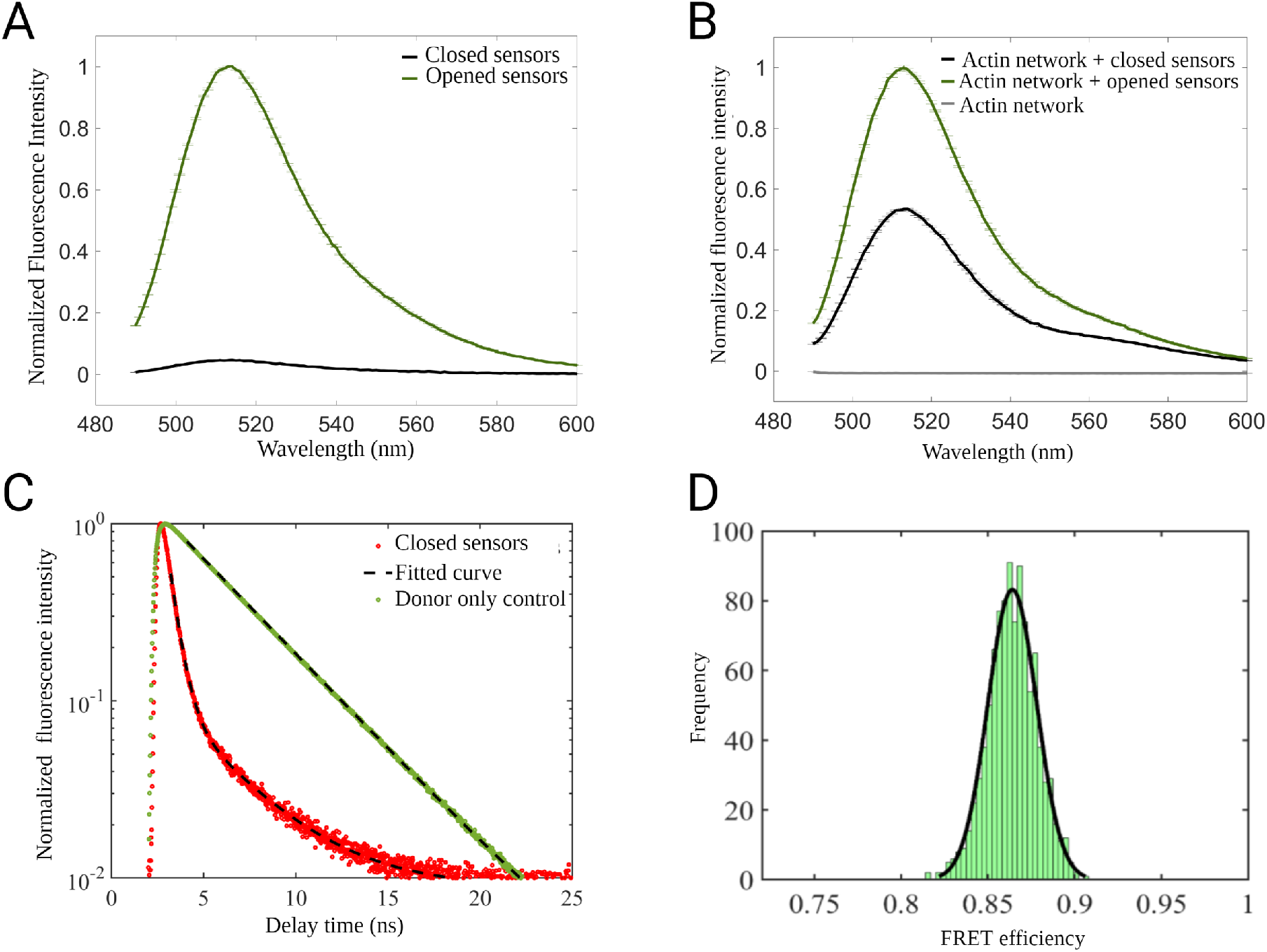
Bulk fluorescence analysis of DNA sensors. Measurements were performed with a commercial spectrometer recording emission spectra at an excitation wavelength of 488 nm. **A:** Opened sensors (with C strand) in solution show a stronger fluorescence signal (green curve) than closed sensors (black curve). **B:** When attached to an actin network, the fluorescence intensity of closed sensors (black curve) is less strongly quenched. **C:** Fluorescence lifetime measurements of sensor molecules in aqueous buffer. Representative TCSPC histograms and fits obtained for closed (red) and donor-only control (green) sensors, respectively. **D:** FRET efficiency calculated by using the measured fluorescence lifetime values of closed and donor only control sensors. Sensor strand stoichiometries are 0.5:1:1 (F:H:Q)

### *In vitro* characterization of DNA sensors by fluorescence lifetime

To avoid possible artifacts inherent to fluorescence intensity measurements, we next measured the donor fluorescence lifetimes (*τ*) and used it to determine the sensor’s FRET efficiency in solution.

FRET efficiency is quantified using the relation

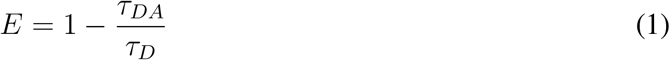

where *E* is the FRET efficiency, and *τ*_*DA*_ and *τ*_*D*_ are the fluorescence lifetimes of donor in presence (closed sensor) and absence (donor-only) of acceptor, respectively. A high FRET efficiency indicates proper assembly of the closed sensor (Fig. 1). All fluorescence lifetime measurements were done by time-correlated single-photon counting (TCSPC) with a confocal microscope ^39^.

We performed further measurements in buffer suitable for *in vitro* actin networks (for buffer composition see Table 3, supporting information). We also optimized the stoichiometry of F, H and Q strands to obtain maximal quenching of fluorescence emission. Fig. 2C presents TCSPC histograms measured for closed sensors and donor-only controls. Fitted fluorescence lifetime values of 0.5 ± 0.1 ns for the closed sensors indicate strong fluorescence quenching, compared with lifetime values of 3.7 ± 0.1 ns for donor-only controls. From the measured lifetime of the closed sensor, we infer a FRET efficiency of 86.0 ± 3.2 % confirming the correct assembly of the sensors. A long lived component of fluorescence for the closed sensors (3.7 ± 0.1 ns) indicates the existence of a non-quenched sub-population of sensors. Likely explanations are the absence of quencher strands in some of the assembled constructs, or misfolding of the constructs themselves. Measured fluorescence lifetime values of opened sensors in DNA hybridization buffer were 3.81 ± 0.03 ns. This is slightly longer than the fluorescence lifetime of the donor-only controls, most probably due to intra-loop quenching of fluorescence by guanosine via electron transfer in the control ^40, 41^. Additional fluorescence lifetime measurements for opened sensors as well as closed sensors containing F, H and Q strands in various molar ratios are also presented (Table 4, Supporting info). These measurements helped us to optimize sensor performance.

### Fluorescence lifetime imaging microscopy (FLIM) of sensors embedded in actin networks

Next, we tested how well the functionality of sensors is preserved when they are linked to actin filaments in a network. Sensor function was again measured with fluorescence lifetime imaging microscopy (FLIM). We reconstituted two kinds of actin networks *in vitro*, cross-linked by closed sensors and by (C-strand) opened sensors respectively, with an actin concentration of 24 µM and a molar ratio of crosslinker to actin-monomer concentration *R* = 0.1. FLIM images were recorded for areas of ∼ 40 µm × 40 µm from six adjacent *z*-planes with a spacing of 1 *µ*m. Figure 3A illustrates the experimental scheme of FLIM on actin networks, cross-linked with sensor constructs. TCSPC curves for Alexa488-tagged sensors inside actin networks are shown in Fig. 3B. Closed sensors in actin networks exhibited a much more rapid fluorescence decay than the ones containing opened sensors. The short (quenched) lifetime component was found to be 1.2 ± 0.1 ns and remained roughly constant across all *z*-planes. However, we also observed also a long lifetime (non-quenched) component of 3.7 ± 0.1 ns. Since the findings are consistent with fluorescence lifetime measurements in ‘actin buffer’, the existence of a non-quenched sub-population of sensors is likely due to binding of actin. Fig. 3C shows distributions of fluorescence lifetime values for closed and opened sensors inside actin networks.

**Figure 3:**
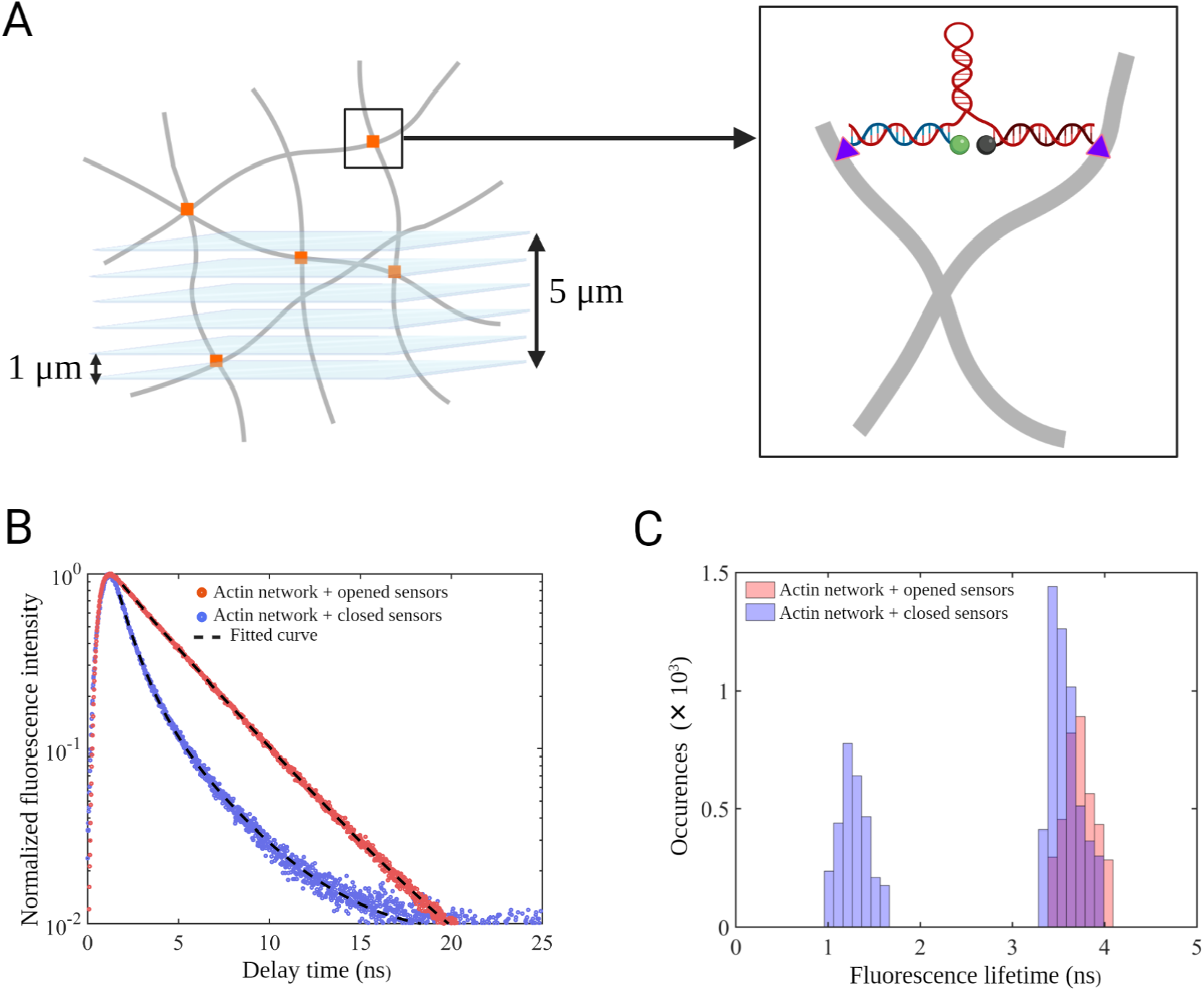
Fluorescence lifetime imaging microscopy (FLIM) of *in vitro* actin networks cross-linked by DNA sensors. **A:** Experimental scheme of *z*-stack FLIM in actin networks. FLIM scans were recorded in different focal planes separated by 1 µm and up to a maximum distance of 5 µm above the glass surface. A zoomed-in schematic of DNA sensors attached to actin filaments is shown on the right. **B:** TCSPC histograms obtained from actin network crosslinked with closed and opened sensors (*R* = 0.1). **C:** Fluorescence lifetime distributions for closed and opened sensors attached to actin networks. From a 2-component fit to the decay, a bimodal lifetime distribution (blue) corresponding to the short and long lifetime components was obtained for closed sensors (1.16 ± 0.08 ns and 3.67 ± 0.06 ns), while the opened sensors exhibit a lifetime of 3.84 ± 0.06 ns (red) while crosslinked to actin network. *R* = c_*crosslinker*_/c_*actin*_.

### Visco-elastic properties of actin networks crosslinked by DNA sensors

Crosslinking of the entangled actin networks by the sensor constructs is expected to strongly affect the viscoelastic properties of the networks and can thus serve as a test for the efficiency of the sensor in linking different filaments as opposed to binding to the same filament. To quantitatively evaluate the viscoelastic properties of sensor-crosslinked actin networks, we measured complex shear moduli *G*(*ω*) of such networks in frequency-sweep experiments between 0.01 and 1 Hz at a strain amplitude of 1% in a rheometer (MCR 501, Anton Paar). In this frequency range we expect elastic plateau behavior with the the real part of *G*(*ω*), the storage modulus *G*′ dominating the imaginary part, the viscous modulus *G*” (Fig. 4 B & Fig. S1 B, C). Overall, sensor-crosslinked networks were substantially more rigid than entangled actin at the same concentration of 24 *µ*M actin. Entangled actin networks showed an elastic plateau at low frequencies with a storage modulus *G*′ = 0.4 Pa, consistent with published studies ^42, 43 44^. Sensor-crosslinked networks displayed plateaus with *G*′ ranging from 0.5 - 1.2 Pa (Fig. 4A & Fig. S1 A), depending on sensor concentration (given by *R* = crosslinker concentration (DNA sensors) / actin concentration in Table 2, supporting information). At the highest sensor concentration that we tested (*R* = 0.2), we found *G*′ = 1.8 Pa for 1:1:1 (F:H:Q) networks (data not shown) and *G*′ = 1.5 Pa for 0.5:1:1 (F:H:Q) networks (Fig. S1 A). At high sensor concentrations (*R* = 0.1 and *R* = 0.2), the network elasticity increased slowly over time and had not reached a steady-state shear modulus after *>*1hr (see Fig. 4A for *R* = 0.1 & Fig. S1 A for *R* = 0.2). Networks with lower sensor concentrations (*R* = 0.005, *R* = 0.01 and *R* = 0.02) had stabilized after ∼2000 s (Fig. 4A & Fig. S1 A). Both the significant increase in plateau modulus and the slow maturation of the crosslinked actin networks towards a steady state is consistent with the reported behavior of actin networks crosslinked by simple double-stranded DNA tethers without sensor functions ^45^.

**Figure 4:**
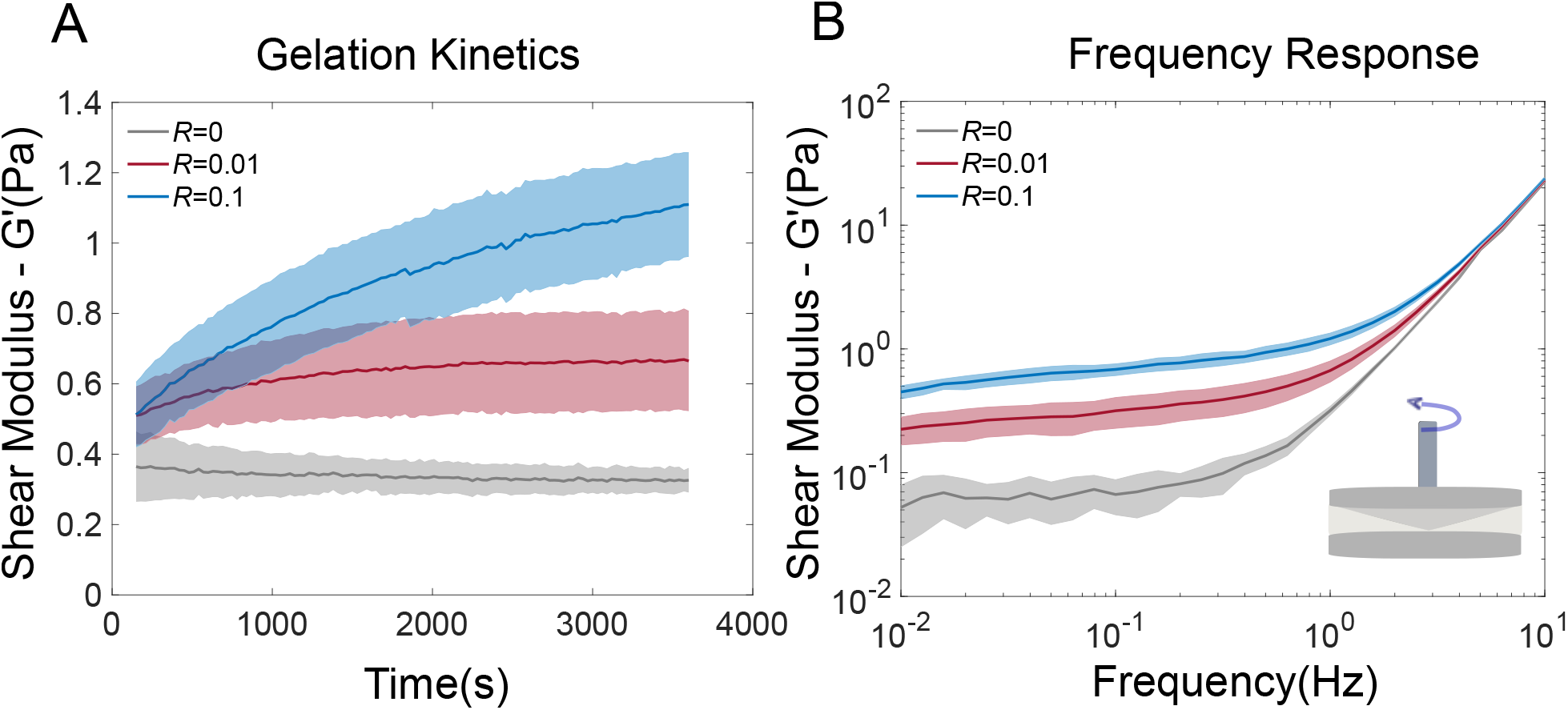
Rheology of actin-DNA sensor networks. **A:** The effect of crosslinking of DNA sensors to actin filaments is observed via an increase in elastic shear modulus (G’, blue and red curves). At a high sensor concentrations (*R* = 0.1), the network become stiffer (increased G’), indicating the formation of a well cross-linked network. **B:** Frequency response of networks in the linear deformation limit at 1% strain. Cross-linked networks do not exhibit any specific frequency-dependent behavior across the probed frequencies. *R* = 0.01 indicates a network with low concentration of sensors, *R* = 0.1 is a network with high sensor concentration, and *R* = 0 indicates an entangled actin network (no sensors). *R* = c_*crosslinker*_/c_*actin*_. Solid lines represent mean values and shaded areas are the standard error of mean.

The changes in structure of sensor-crosslinked actin networks that we observed in a confocal microscope corresponded to changes of the measured elasticity. At low sensor concentrations (*R* = 0.005, *R* = 0.01, *R* = 0.02), networks appeared isotropically crosslinked (Fig. 5 A,B,C,D). At higher sensor concentrations, *R* = 0.1 and *R* = 0.2 (Fig. 5 E,F), the networks became inhomogeneous, with denser actin bundles co-existing with a homogeneous network background. These bundles were always observed throughout the samples (Fig. S2), proving that they were not surface artifacts. Such composite networks including isolated bundles were also observed by Lorenz *et al*. ^45^ in their study of ds-DNA-actin networks. The slow approach to a mechanical steady state (Fig. 4A, *R* = 0.1) observed in the rheology experiments likely reflects ongoing formation of bundles that would eventually percolate. Bundles in our networks are observed only at high concentrations of sensors and thus cannot come from phase separation driven by solvent conditions, i.e. changes in ion concentrations ^46^. Bundles are also unlikely to be driven by depletion forces due to DNA. Entropy-driven bundle formation should result in a loss of connectivity/entanglements in actin ^47^. This is not the case for our sensor-crosslinked actin networks since their elasticity remains high, indicating good connectivity between sensors and actin.

**Figure 5:**
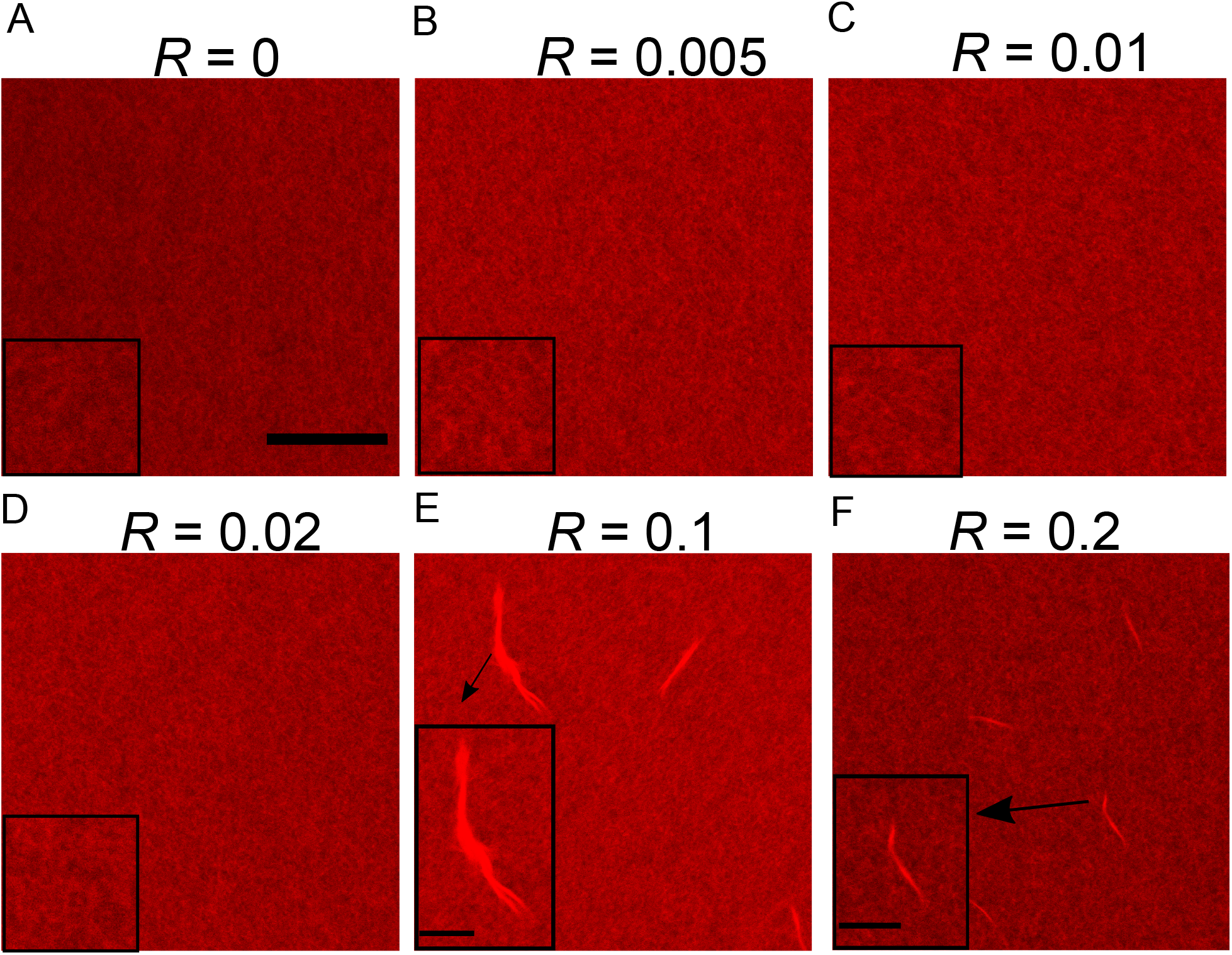
Morphology of DNA-sensor-crosslinked actin networks. Confocal laser scan images of **A:** entangled actin (*R* = 0) and **B-F:** DNA-sensor-crosslinked networks (from *R* = 0.01 to *R* = 0.2). An isotropically crosslinked network is observed for all lower *R*-values (*R* = 0.005, *R* = 0.01, *R* = 0.02) which is not visibly different from an entangled actin network (*R*=0). At higher sensor concentrations (*R* = 0.1, *R* = 0.2), composite network structures arise which have the appearance of bundles embedded in a crosslinked network. *R* = c_*crosslinker*_/c_*actin*_. Scale bar: 30 µm. Inset scale bar: 10 µm. Insets of A,B,C,D are zoomed images. Insets of E and F are zoomed images for the bundled indicated. Actin is fluorescently labeled with Atto 647N-Phalloidin.

### Sensor characterization inside living cells

Last, we investigated the suitability of our sensors for live cell measurements. To this end, HeLa cells were transfected with HaloTag®-RFP-LifeAct, followed by microinjection of a 50 nM solution of closed DNA sensors (see Fig. S3 for details). Figure 6A shows confocal micrographs of a HeLa cell in both sensor (left) and RFP (actin, middle panel) spectral channels. The images show excellent co-localization of sensors and actin stress fibers (Fig. 6A right panel, Fig. S4). We observed an average fluorescence lifetime of 2.2 ± 0.5 ns for dye-tagged DNA sensors inside living cells as seen in Fig. 6B, Fig. 6C and Fig. S5. These findings confirm that the MFS introduced here is applicable for FLIM imaging inside living cells which opens up possibilities of direct quantification of cellular forces.

**Figure 6:**
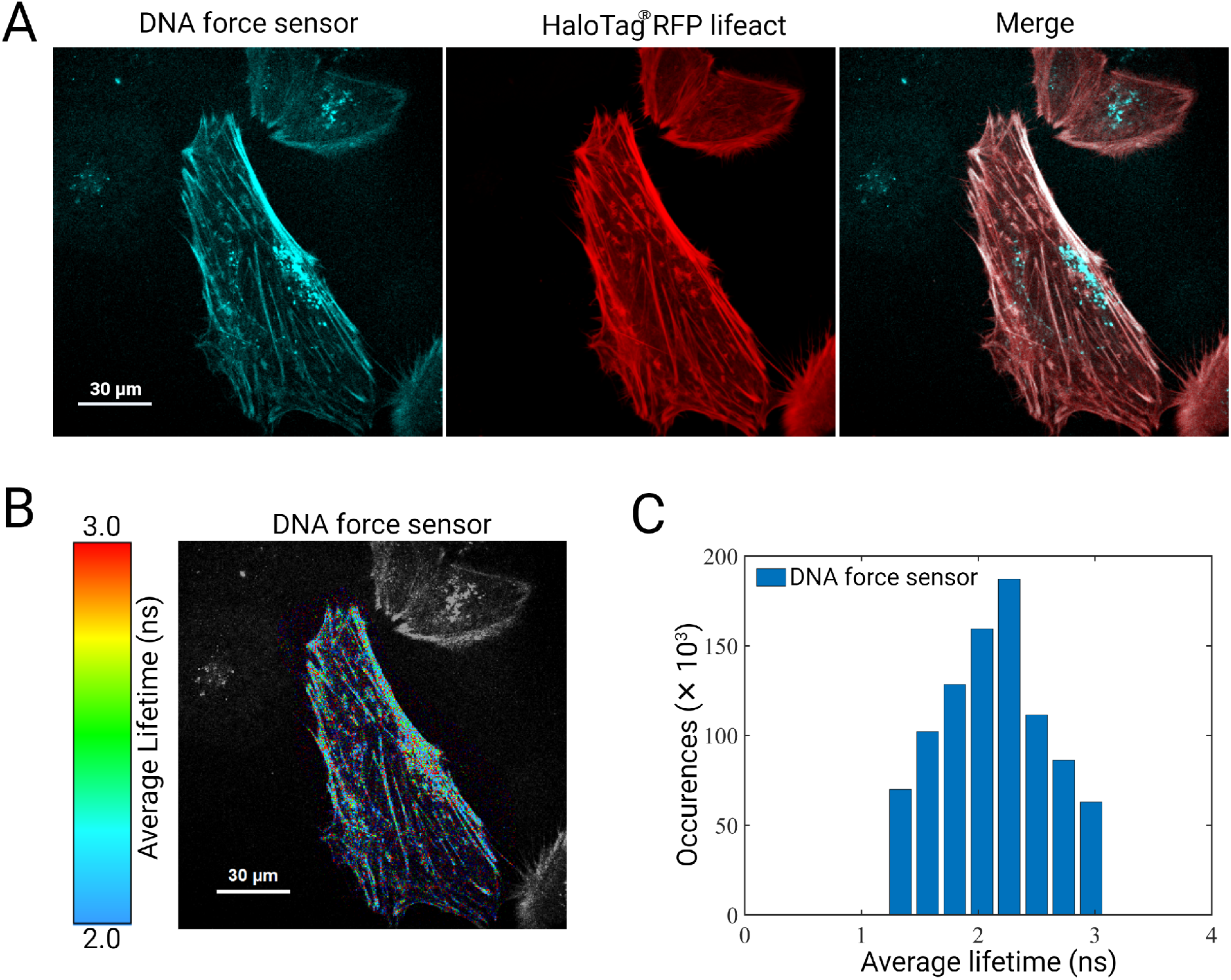
HeLa cells microinjected with closed DNA sensors. **A:** Confocal micrographs of a cell false-coloured for fluorescence intensity. Images corresponding to Alexa 488-tagged closed DNA sensor recorded in the DNA force sensor spectral channel (left), the same cell in RFP spectral channel (middle) and the overlay between both channels (right) confirms co-localization between DNA-sensors and actin. **B:** FLIM image of the same cell as in **A** is illustrated for the DNA force sensor spectral channel. **C:** Bar histogram showing fitted fluorescence lifetimes obtained from **B**. An average fluorescence lifetime of 2.2 ± 0.5 ns was calculated for DNA force sensors.

In this feasibility study we have constructed and tested a DNA-based molecular force sensor, designed to attach to and crosslink actin filaments in the cytoskeleton of cells. Our sensors are meant to sample cell-internal stresses that would not necessarily be transmitted to the cell environment. The sensors use FRET between a synthetic organic dye and a quencher that can be read using fluorescence intensity or fluorescence lifetime imaging. Sensors can be used in *in vitro* experiments or in cells. They can be directly attached to any cellular component (here we targeted actin) in a flexible manner with the help of binding proteins or fragments of such using the HaloTag® system. We tested the sensors’ binding characteristics in actin networks, a major component of the cytoskeleton of cells. Quenching efficiency was 86.0 ± 3.2 %, which is comparable to other reported DNA force sensors which, in contrast to ours, were designed for surface attachment^29, 36^. Our 8 bp hairpin DNA sensors can easily be modified by changing hairpin length and GC content to probe different force ranges ^32–36^. In our case of actin crosslinking sensors, various binding tools ^48^, or mutant ABP’s can be employed. Binding proteins or fragments have broadly varying off-rates ^49, 50^, which would endow the sensors with a temporal high-pass filter function: only force changes that occur faster than the inverse off-rate will be detected. The design of our sensor is a first step towards the study of force transmission across cellular biopolymer networks. A next major challenge will be to explore the effects of attachment geometry. In contrast to sensors that have been deployed between cells and substrates or were embedded in more or less 1-D load bearing structures such as focal adhesions, the network sensors will likely bind in very inhomogeneous directions and configurations, experiencing broadly varying force on the molecular scale in deforming networks. The technology we introduced here will open the door towards sampling these forces all-optically and non-invasively in complex environments such as cells and tissues.

## Methods

### Sensor sequence design and chemical modifications

Sensor sequences (see Supporting Table 1) were designed on the NUPACK website ^51^, and DNA oligomers were purchased from Integrated DNA Technologies (Leuven, Belgium). Lyophilized powders were reconstituted to 100 or 500 µM stock concentrations by adding an appropriate volume of DNA hybridization buffer (see Supporting Table 3). Sensors were further chemically modified at the 3’ end of the F strand and the 5’ end of the Q strand (see Table 1, supporting info). These strands were purchased with thiol groups incorporated at the respective ends (see supporting info Table 1). Both strands were separately modified. Between 50 µl and 100 µl of 30 µM solutions of these strands were prepared by dilution in DNA hybridization buffers and reduced with 2 mM TCEP (Tris(2-Carboxyethyl)phosphine hydrochloride, Sigma-Aldrich, Germany) by incubating at room temperature for 60 minutes. Next, iodoacetamide (O_4_) HaloTag® ligand (Promega, Madison, USA) was added to the above solution to a final concentration of 20 µM and incubated for 90 min.

### Assembly of DNA sensors

A volume of 50 µl of a solution of closed sensors or donor-only controls was prepared by adding modified F strand (to 5 µM), modified Q or Q^™^ strand (to 10 µM) and H strand (to 10 µM final concentration) to DNA buffer. Opened sensors were assembled in the same way, followed by addition of C strand (complementary strand to hairpin loop). The molar concentration of C strand was 10 times larger than that of H strand. The final concentrations of sensors and donor-only controls were varied depending on the experiment (see Table 2) while the ratios were kept constant at 0.5:1:1 (F:H:Q) in all experiments.

### Actin and actin-DNA sensor network preparation

G-actin was prepared from rabbit skeletal muscle according to published protocols ^52–54^. Purified actin was stored in small aliquots at −80°C. Aliquots were thawed freshly prior to experiments. Concentrated 10x polymerization buffer (for actin buffer composition see Supporting Table 3, Polymix 10X, Hypermol) was added to an appropriate volume of water. G-actin was added to reach a final concentration of 24 µM (1 mg/ml). The concentration of actin remained unchanged throughout the experiments. To construct actin-DNA sensor networks, Lifeact-RFP-HaloTag® protein was added to a final concentration of 30 µM. Then modified and assembled sensors or donor-only controls, prepared as described above, were added. The sensor concentrations were varied from 0 µM to 4.7 µM (*R*-ratio, see Supporting Table 2). For imaging experiments, a 10 % volume from a 10 nmol stock solution of Atto 647N, phalloidin (Atto-TEC GmbH, Siegen) was finally added. The solutions were gently pipetted to distribute components and then filled into a microscope chamber.

The LifeAct-RFP-HaloTag® proteins used to attach the sensors to *in vitro* actin networks were bacterially expressed and purified following standard procedures and added to the above described modified F and Q strand solutions. The LifeAct-RFP-HaloTag® covalently binds to the iodoacetamide (O_4_) HaloTag® ligand on the F and Q strands.

### Fluorescence intensity measurements

Fluorescence intensity measurements were performed with an AMINCO-Bowman Series 2 Luminescence Spectrometer (Thermo Electron Scientific Instruments Corporation, Madison, WI 53711, USA). Emission spectra were measured with the following instrument settings: Excitation wavelength - 488 nm, emission wavelength - 520 nm, bandpass - 1, emission scan range 490 - 600 nm.

### Rheology measurements

Rheological measurements were performed with a commerical rheometer (MCR 501 Anton Paar, Austria) in a cone-plate geometry (CP-25, 2°). Polymerization of networks was monitored in time-sweep measurements with an oscillatory shear at 1% strain amplitude, 1 Hz frequency and at a temperature of 23°C. To prevent evaporation during the measurements, sample hydration was maintained by placing wet tissue paper around the measurement plates. The storage and loss moduli of networks were obtained. Viscoelastic properties of the network were determined from frequency-sweep measurements, performed after the time sweeps for all networks. The frequency range was 0.01 Hz to 10 Hz at 1% strain (see Fig. S1 and Fig. S2). Phalloidin was not added to the networks used for rheological experiments.

### Confocal imaging of networks

Microscope chambers were constructed from KOH-cleaned microscope slides (631-1550, VWR, Germany) and cover slips (No. 1.5, 24×24 mm, VWR, Germany), using two strips (3 mm wide) of optically clear double-stick adhesive tape (50 µm thick, 3M™, #8212, USA). 50 µl of actin or actin-DNA sensor solution was carefully deposited in the middle of the chamber before closing it. Solutions spread when gently placing the coverslip on top. Care was taken to avoid air bubbles in the chamber. The chambers were then immediately sealed with VALAP (vaseline, lanolin, and paraffin sealant mix) and wrapped with aluminum foil to prevent photobleaching during polymerization. Imaging was performed after the completion of polymerization (1 hour). Networks were imaged with a Leica TCS SP5 confocal microscope (Leica Microsystems CMS GmbH, Mannheim, Germany) to probe the morphology of the resulting networks. We recorded confocal scans across the entire depth of the networks (*z*-scans). A whitelight laser (specs) was used for illumination with 20 % maximum intensity for the sensor channel (excitation wavelength - 488 nm) and 22 % maximum intensity for the actin network imaging channel (excitation wavelength - 647 nm). An xyz image acquisition (1024 × 1024 pixels) was done with a bidirectional scan with each line scanned for 16 times (Line average = 16) at 700 Hz speed. The resulting field of view was 100 µm x 100 µm. A zoom factor of 2.5 was used. A pinhole size of 1 Airy unit (95.4 µm) was used, and *z*-stacks were obtained with a 3 µm step size.

### Fluorescence lifetime measurements in aqueous buffer

A home-built confocal microscope capable of fluorescence lifetime measurements was used. Excitation was done with a linearly polarized pulsed diode laser (*λ* = 485 nm, pulse duration 50 ps FWHM, LDH-P-C-485B, PicoQuant) equipped with a clean-up filter (Brightline FF01-480/17, Semrock). Light of this laser was pulsed at a repetition rate of 40 MHz with a multi-channel picosecond laser driver (PDL 828, “Sepia II”, PicoQuant). The laser beam is coupled into a polarization-maintaining single-mode fiber (PMC-400-4.2-NA010-3-APC-250V, Schäfter and Kirchhoff GmbH). At the fiber output, the light is colli-mated and reflected by a dichroic mirror (FITC/TRITC Chroma Technology) into the objective lens of the microscope (UPLSAPO 60x water, 1.2 NA, Olympus). The same water-immersion objective is used to collect fluorescence from the sample. A long-pass filter (BLP01-488R-25, Semrock) is used to block back-scattered light from the laser. The emission light is focused into a pinhole of 100 µm diameter, collimated again, refocused onto a single-photon avalanche photo-diode (SPCM - CD 3516 H, Excelitas Technologies GmbH & Co. KG). A multi-channel picosecond event timer (HydraHarp 400, PicoQuant) records the detected photons from the detector with an absolute temporal resolution of 16 ps. For lifetime measurements, a droplet of 30 µl of DNA sensors dissolved in DNA hybridization buffer (see table 3 for buffer composition) was placed on a glass coverslip (24 mm × 24 mm, thickness 170 µm). The laser beam was focused 30 µm inside the droplet, and data acquisition was started. TCSPC histograms were computed from the recorded photons and mono-exponential (opened sensors and donor only controls) and bi-exponential (for closed sensors) decay functions were fitted to the tails of the histograms (0.5 ns after the maximum) using a maximum-likelihood procedure as described elsewhere ^55^. We also performed fluorescence lifetime measurements on DNA sensors dissolved in an aqueous buffer suitable for actin networks. A further goal of *in vitro* fluorescence lifetime measurements was to screen for an optimal molar ratio between the individual strands such that sufficient F strands are present for cross-linking actin filaments. It was equally important to obtain a molar ratio which ensures maximal quenching of fluorescence lifetimes. For meeting both the criteria, we varied the molar concentration of the F strand in closed sensor and determined fluorescence lifetimes for four different molar ratios, 2:2:2, 1:2:2, 0.5:2:2 and 0.25:2:2 (F:H:Q). Table 4 (Supporting information) lists the fluorescence lifetime values as obtained from TCSPC for all the different constructs. These values suggest that a molar ratio of 0.5:2:2 contains sufficient F strands to meet both criteria.

### Introduction of sensors into living cells

Introduction of sensors into living cells was performed in a two-step procedure (see Fig. S3). HeLa cells (ACC 173, Leibniz Institute DMSZ, Braunschweig, Germany) were cultivated to confluency in T75 flasks and passaged as follows. They were rinsed with 10 ml of PBS (phosphate buffer saline), trypsinized for 3 mins with 5 ml of an EDTA/Trypsin solution (0.05 %, 59417C, Gibco, Thermo Fisher), and diluted with low glucose DMEM (Dulbecco’s modified Eagle’s medium, SigmaAldrich, St.Louis, MO, USA) containing 10 % fetal bovine serum (FBS) (# F0244, Sigma-Aldrich, heat-inactivated (30 min, 56 °C) and 1 % penicillin-streptomycin (# 17-602E, Lonza, Basel, Switzerland). The cell solution was then centrifuged for 5 min at 1000 rpm, and the resulting pellet was resuspended in 1000 µl DMEM. 40,000 cells were plated on ibidi µ-dishes (#81166, ibidi, Germany). 48 hrs post cell seeding in the ibidi dishes, cells were transfected with the lifeact-RFP-HaloTag® plasmid using lipofectamine 3000 reagent (L3000001, Thermo Fisher Scientific, Germany). After ∼24 h expression, small volumes of a 50 nM DNA sensor solution in Phenol red free medium was microinjected into the cells using a Transjector 5246 (Eppendorf, Hamburg, Ger, 5246 01084) in combination with Femtotips (Eppendorf, Hamburg, Ger, 930000035). Injection was performed close to the nucleus in 3 s pulses.

### FLIM measurements in live cells

Lifetime measurements were performed within 1 hour after microinjection of the sensor construct into HeLa cells. The imaging medium consisted of: Phenolred free medium (DMEM,Gibco, 1X, 11880-028, Life Technologies, UK), with 10 % fetal bovine serum (FBS), heat-inactivated (30 min, 56 °C),(# F0244, Sigma-Aldrich) and 1 % penicillinstreptomycin (# 17-602E, Lonza, Basel, Switzerland). FLIM measurements were performed using the same custom-built confocal microscope described above. Briefly, the laser beam was focused on single cells, and FLIM scans were recorded from multiple regions of interest. TCSPC histograms of each pixel were computed and then fitted using a bi-exponential decay function. An average lifetime value was calculated for each pixel, weighing the lifetime components with their respective fluorescence photons (as used for TCSPC fitting). The obtained values of intensity-weighted average lifetimes are color-coded in the FLIM images in Fig. S4 of supporting information and Fig. 6 in the main text.

## Data availability

All data that support the findings described in this study are available within the manuscript and the related Supporting Information, and from the corresponding authors upon request.

## Acknowledgements

The authors thank Fred Wouters for insightful discussions. This work was financially supported by the Deutsche Forschungsgemeinschaft (DFG, German Research Foundation) via project A03 (F.R. and C.F.S.) of the Collaborative Research Center 755 (SFB 755). J.E. was also supported by the Deutsche Forschungsgemeinschaft (DFG, German Research Foundation) under Germany’s Excellence Strategy - EXC 2067/1-390729940.

## Author contributions

F.R. and C.F.S. conceived the study; F.R. and C.F.S., A.T. and M.P. designed the DNA constructs, M.P.performed preliminary experiments. C.J. (sensor preparation, bulk fluorescence, networks imaging, rheology, cell culture) and A.G. (FLIM imaging) performed the experiments reported here with the help of L.H. for cell experiments (microinjection). J.E. supervised the FLIM experiments, F.R. and C.F.S supervised all other experiments, C.J., A.G., J. E., F.R. and C.F.S. wrote the manuscript.

