## supporting figures S1,S2,S3,S4,S5 and tables 1, 2, 3, 4 for "A DNA-based optical force sensor for live-cell applications"

Table 1: DNA oligos sequence

| DNA oligo strands | Sequence |
|---|---|
| F | 3' Thiol C3 GCGGGACTTTCGTGCGTCGC Alexa 488 5' |
| Q | 3' Iowa blackFQ CGCGCCCGTGCGCCGAACGC C6 Thiol 5' |
| H | GCGAACCG GAGAGTGTTAGAGACA CGGTTCGC |
| C | CTCTCACAATCTCTGTCGGTTCGC |

Table 2: DNA sensor crosslinking concentration

| Network | $R$ | Sensor concentration ($\mu$M) |
|---|---|---|
| Actin[a] | 0 | 0 |
| Actin[a]+DNA sensor | 0.005 | 0.12 |
| | 0.01 | 0.24 |
| | 0.02 | 0.48 |
| | 0.1 | 2.4 |
| | 0.2 | 4.7 |

[a] A fixed actin concentration of 23.81 $\mu$M was always used;

Table 3: Buffer compositions

| Buffer types | Composition |
|---|---|
| DNA hybridization | 50 mM NaCl, 10mM $MgCl_2$, 1x PBS |
| Actin buffer | 1M KCl, 0.1M imidazole pH 7.4, 10mM ATP, 20mM $MgCl_2$ |

| Closed sensor | $\tau$ [ns] DNA hybridization buffer | $\tau$ [ns] actin buffer |
|---|---|---|
| **F:H:Q (2:2:2)** | 0.62 ± 0.04 [0.25] | 0.80 ± 0.08 [0.22] |
| | 3.70 ± 0.07 [0.75] | 3.58 ± 0.07 [0.78] |
| **F:H:Q (1:2:2)** | 0.53 ± 0.04 [0.36] | 0.59 ± 0.02 [0.25] |
| | 3.70 ± 0.06 [0.64] | 3.58 ± 0.07 [0.75] |
| **F:H:Q (0.5:2:2)** | 0.53 ± 0.08 [0.38] | 0.59 ± 0.04 [0.29] |
| | 3.70 ± 0.07 [0.62] | 3.58 ± 0.05 [0.71] |
| **F:H:Q (0.25:2:2)** | 0.51 ± 0.08 [0.43] | 0.61 ± 0.05 [0.29] |
| | 3.70 ± 0.08 [0.57] | 3.58 ± 0.05 [0.71] |

Table 4: Fluorescence lifetime values (in ns) and their relative fit amplitudes

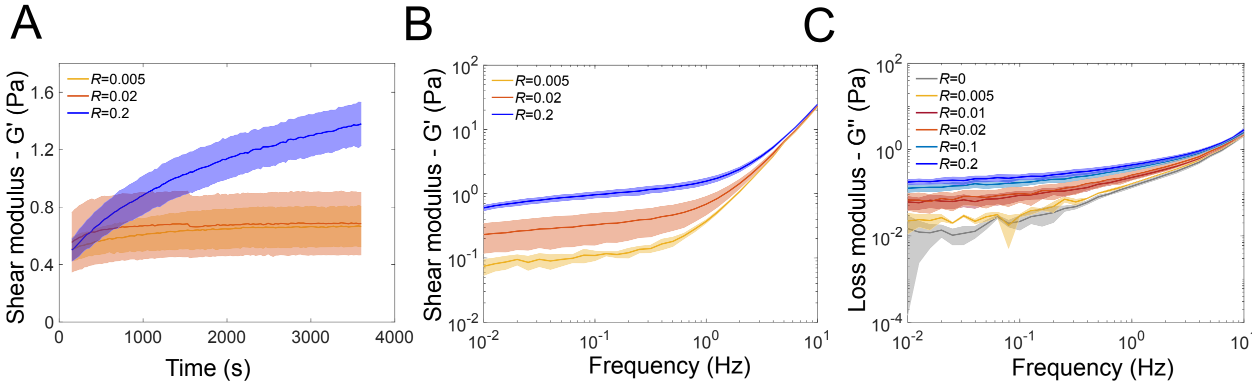


Figure 1: Gelation kinetics and frequency response of actin, actin-DNA sensors networks at different sensor concentrations. A,B,C. Gelation kinetics for sensor concentrations ($R$ = 0.005, 0.02, 0.2) to show the evidence of a crosslinked network. Frequency response from these networks also do not exhibit any distinctive frequency dependent behavior.Solid lines represent mean values and the shaded area the standard error of mean.

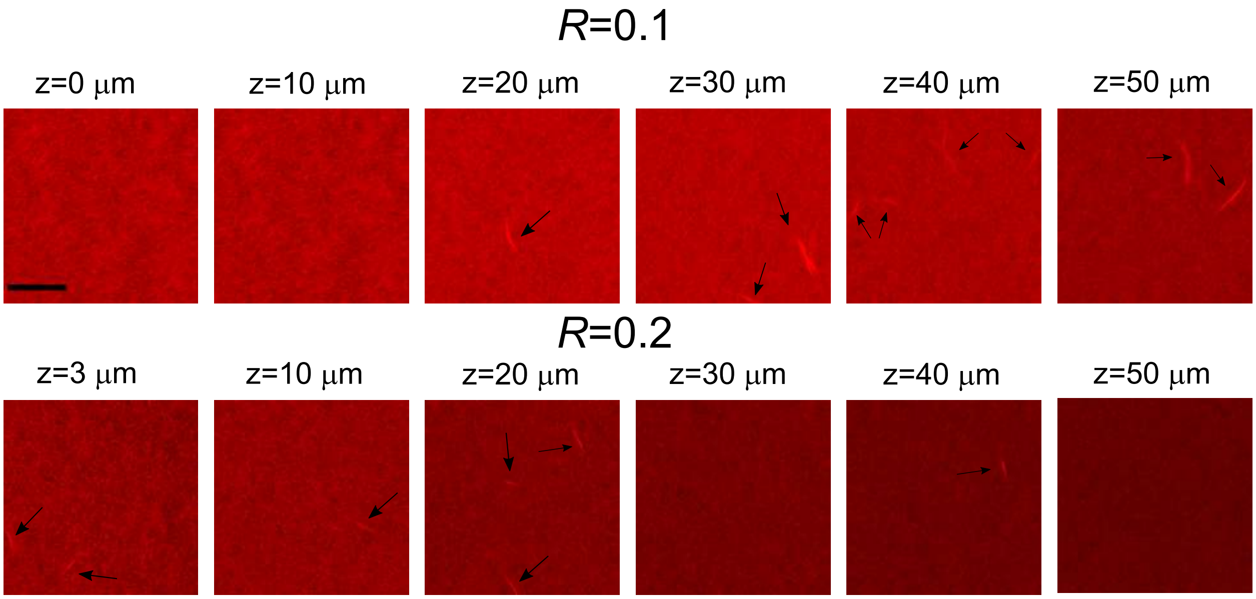


Figure 2: Microstructure of $R$ = 0.1 & $R$ = 0.2 actin-DNA sensor networks at varying z-heights in the network. Actin-sensor networks at high sensor concentrations ($R$ = 0.1, 0.2) were imaged in a confocal z-scan which shows the existence of bundles across different z-positions. They appeared mostly in the middle of the network and were not present at the boundary surfaces of the coverslip and the microscopic slide. (Scale bar: 30 $\mu$m). Actin is labeled for fluorescence with Atto 647N.

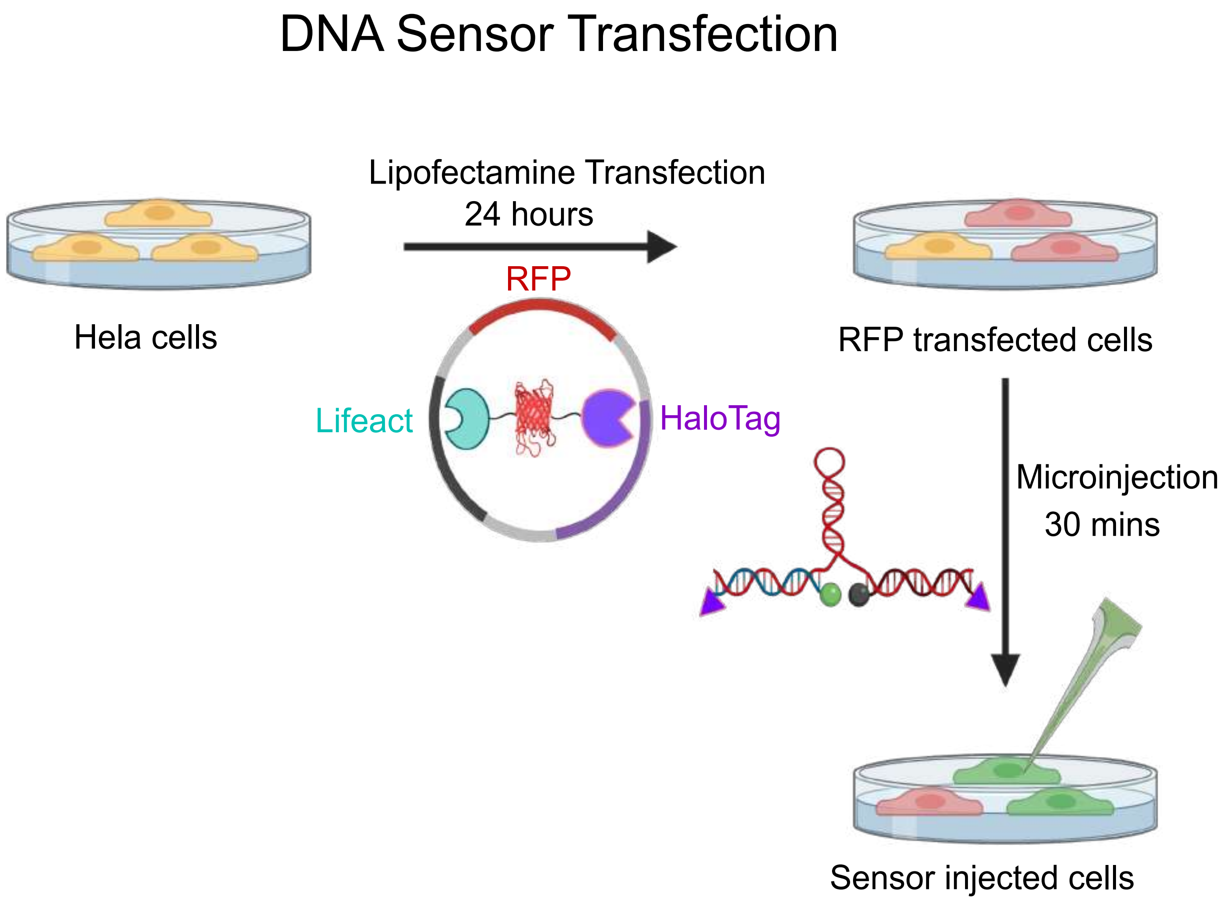


Figure 3: A schematic representation of the transfection protocol of DNA sensors. DNA sensors were transfected into Hela cells in 2 steps. The lifeact-RFP-HaloTag® plasmid was transfected into Hela cells with lipofectamine 3000. 24h later, a 50 nM solution of DNA sensors that were first modified, were assembled *in vitro* and microinjected near the nucleus. (Scheme is not drawn to scale.)

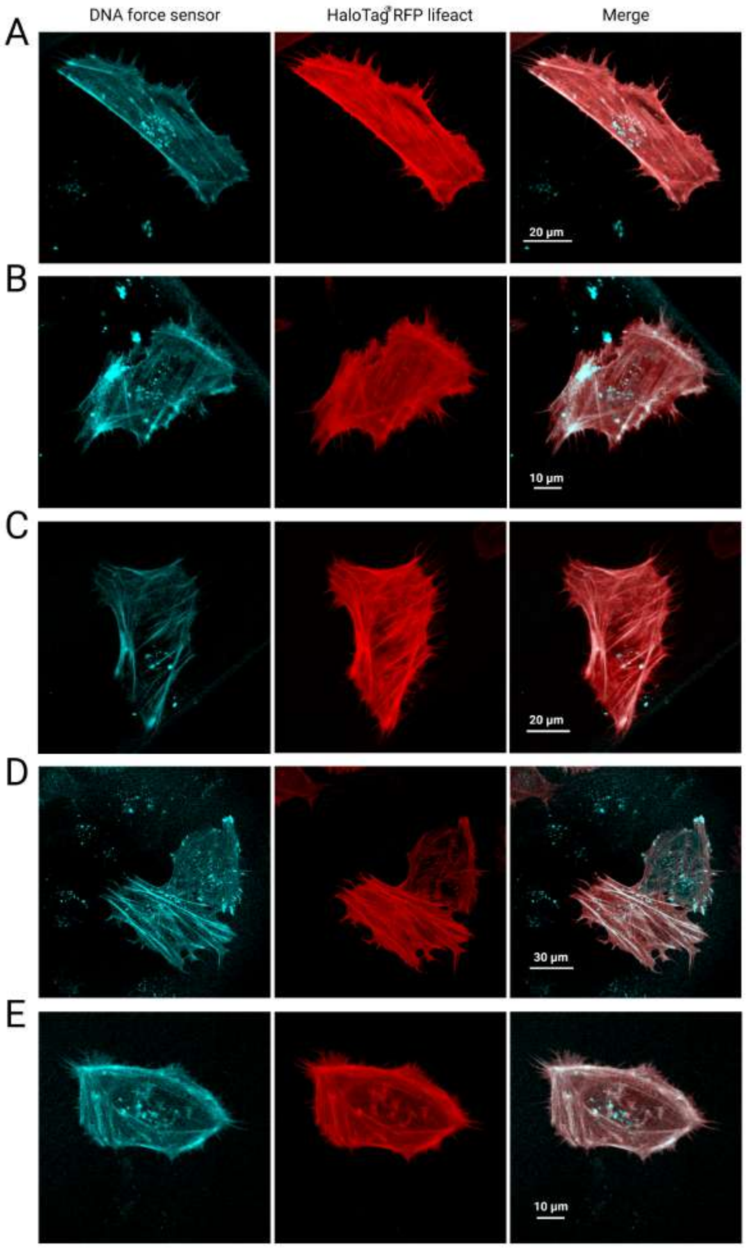


Figure 4: **A** - **E** HeLa cells microinjected with closed DNA sensors. Confocal micrographs of cells false-coloured for fluorescence intensity. Images corresponding to Alexa 488-tagged closed DNA sensor recorded in the DNA force sensor spectral channel (left), the same cell in RFP spectral channel (middle) and the overlay between both channels (right) confirms co-localization between DNA-sensors and actin.

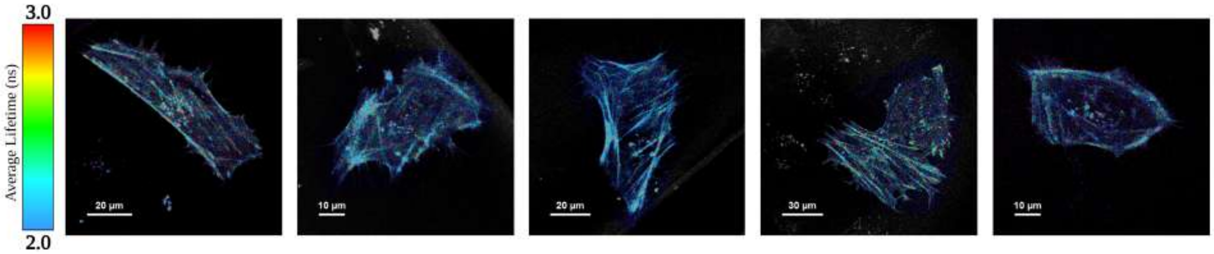


Figure 5: FLIM images of HeLa cells microinjected with DNA sensors.
